# Supramolecular architecture of the ER-mitochondria encounter structure in its native environment

**DOI:** 10.1101/2022.04.12.488000

**Authors:** Michael R. Wozny, Andrea Di Luca, Dustin R. Morado, Andrea Picco, Patrick C. Hoffmann, Elizabeth A. Miller, Stefano Vanni, Wanda Kukulski

## Abstract

The endoplasmic reticulum and mitochondria are main hubs of eukaryotic membrane biogenesis which rely on lipid exchange via membrane contact sites, but the underpinning mechanisms remain poorly understood. In yeast, tethering and lipid transfer between the two organelles is mediated by the ER-mitochondria encounter structure ERMES, a four-subunit complex of unclear stoichiometry and architecture. We determined the molecular organization of ERMES within cells using integrative structural biology, combining quantitative live-imaging, cryo-correlative microscopy, subtomogram averaging and molecular modeling. ERMES assembles into approximately 25 discrete bridge-like complexes distributed irregularly across a contact site. Each bridge consists of three lipid-binding SMP domains arranged in zig-zag fashion. Our molecular model of ERMES reveals an unconventional restrained pathway for lipids. These findings resolve a supramolecular architecture controlling interorganelle lipid fluxes.

## Introduction

Lipids are fundamental constituents of eukaryotic cells and are mainly synthesized in the endoplasmic reticulum (ER). The ER supplies lipids to other organelles for membrane homeostasis and expansion. Although lipids and proteins traffic via vesicles from the ER to outlying organelles, the bulk of cellular lipid flux occurs through membrane contact sites (MCS)(1), which are regions where two organelles are physically apposed (2). For lipid flux between the ER and mitochondria, MCS are the only conduit (3-5). In yeast, ER-mitochondrial MCS are mediated exclusively by the ER-mitochondria encounter structure (ERMES), which is necessary for respiratory function, mitochondrial morphology, and maintenance of the mitochondrial genome (6, 7). ERMES mediates the transport of phospholipids between ER and mitochondria (8, 9), possibly underpinning its roles in mitochondrial function. ERMES is composed of four core components, Mmm1, Mdm12, Mdm34 and Mdm10, which have molecular weights between 31 and 56 kDa (6). Three of them share conserved synaptotagmin-like mitochondrial-lipid-binding (SMP) domains, characteristic of many MCS lipid transfer proteins in yeast and mammals (10-12). Although structural information on SMP domains is available, including for Mmm1 and Mdm12, it remains unknown how SMP domains arrange between two organelles to drive inter-organelle lipid transfer (13-17). Specifically, whether ERMES subunits arrange into stoichiometric complexes that form lipid shuttles or continuous conduits is unclear.

## Results and Discussion

To reveal the supramolecular structure of ERMES-mediated MCS in budding yeast, determine the *in situ* organization of ERMES SMP domains and obtain insights into the mechanism of lipid transfer, we used an integrative structural biology approach. When observed by fluorescence microscopy (FM), the three SMP-domain containing ERMES components Mmm1, Mdm12 and Mdm34 organize as diffraction limited puncta that correspond to ER-mitochondria MCS (Figure 1A)(6). We first determined the number of ERMES components per puncta and their ratio by quantitative live cell FM (18, 19). Fluorescence intensity measurements provided us with an estimate of the absolute number of molecules per MCS of the three ERMES components (Figure 1B). The number of Mmm1 and Mdm34 molecules matched closely (25-27 and 26-29 molecules/punctum, respectively), and was also similar for Mdm12 (19-22 molecules/punctum) (Figure 1C). We thus propose that ER-mitochondria MCS consist of similar protein copy numbers of the three SMP-domain containing ERMES components. The lower abundance of Mdm12 could reflect its assembly properties, as Mdm12 is cytosolic and its targeting to MCS might require its interaction partners Mdm34 and Mmm1 to be present (14, 20). In contrast, Mmm1 is anchored to the ER by a transmembrane domain, and Mdm34 associates tightly with the outer mitochondrial membrane (OMM) protein Mdm10 (20, 21). Nevertheless, considering that the SMP domains of Mmm1 and Mdm12 form a stable equimolar complex *in vitro* (16), our finding supports a model of ERMES assembling as a protein complex with equimolar stoichiometry between Mmm1, Mdm12 and Mdm34.

**Figure 1.**
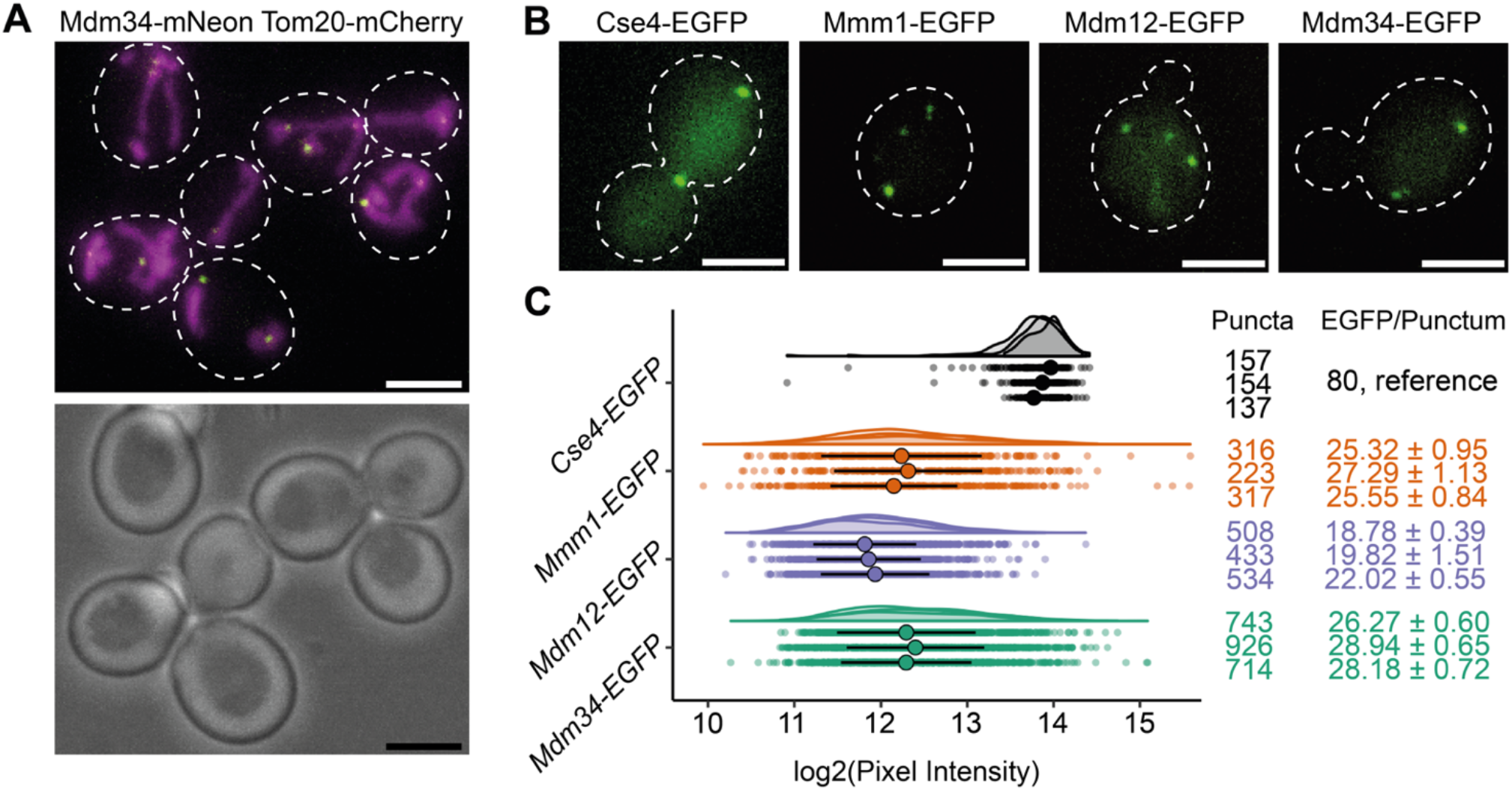
The number of molecules of ERMES components per MCS. **A:** Live cell imaging of budding yeast cells expressing Tom20-mCherry, marking mitochondria, and Mdm34-mNeonGreen, marking ERMES-mediated MCS. In the fluorescence image (top) white dashed outlines mark cell boundaries according to bright field image (bottom). **B:** Live cell FM of yeast cells expressing either Cse4-EGFP, Mmm1-EGFP, Mdm12-EGFP or Mdm34-EGFP. White dashed outlines mark cell boundaries. Cells expressing the kinetochore protein Cse4-EGFP, of which the number of molecules per diffraction limited spot is known (36) were used as a reference to determine the number of molecules of ERMES components. **C:** Fluorescence intensity quantifications of diffraction-limited puncta of EGFP-tagged Cse4 (grey), Mmm1 (orange), Mdm12 (purple) and Mdm34 (green), represented as dot plots as well as half-violin plots. For each quantification, three experimental repeats are shown. Large dots represent the median, lines the MAD, of each experimental repeat. Left column indicates number of analyzed puncta. Using Cse4-EGFP as reference, fluorescence intensities were transformed into numbers of EGFP molecules/punctum (right column), of which median values with MAD are given. Scale bars are 3 µm.

A stoichiometric ERMES complex is compatible with both proposed functional models: that of a stable conduit bridging the ER to mitochondria, or of a mobile shuttle moving between these two organelles. To determine the mode of lipid transport at ER-mitochondria MCS, we visualized ERMES MCS in vitrified yeast cells using correlative light and electron cryo-microscopy (cryo-CLEM). Vitrified cells were thinned into lamellae using cryo-focused ion beam (FIB) milling (22), then imaged using cryo-FM to determine the presence and position of ERMES puncta, marked by Mdm34-mNeonGreen, within each lamella (Figure 2A). Regions of lamellae with ERMES puncta were then imaged using electron cryo-tomography (cryo-ET). This approach reliably directed data collection to ER-mitochondria MCS. These MCS consisted of diverse morphologies of ER and mitochondria (Supplementary Figure S1). ER membranes in contact with mitochondria did not show preferred curvature: convex (28%), concave (33%), convex and concave in different locations of the same MCS (5%), or nearly flat (33%, N=60 MCS) membranes were all observed. Generally, the ER in the MCS appeared to be pressed against the OMM (Supplementary Figure S1). In some cases, the ER was wrapped around the mitochondrial surface (Supplementary Figure S1A and C); in others, ER tubules passed through holes in mitochondria (Supplementary Figure S1D and E). Occasionally, we found peroxisomes near ER-mitochondria MCS, often in close contact with the ER (Supplementary Figure S1B and E). Notably, in addition to 51 tomograms of ER-mitochondria MCS, 9 tomograms collected at fluorescent ERMES puncta contained no ER-mitochondrial MCS but instead ER-peroxisome MCS (Supplementary Figure S2). These observations are in line with previous findings that ERMES can localize to MCS involving peroxisomes (23, 24).

**Figure 2.**
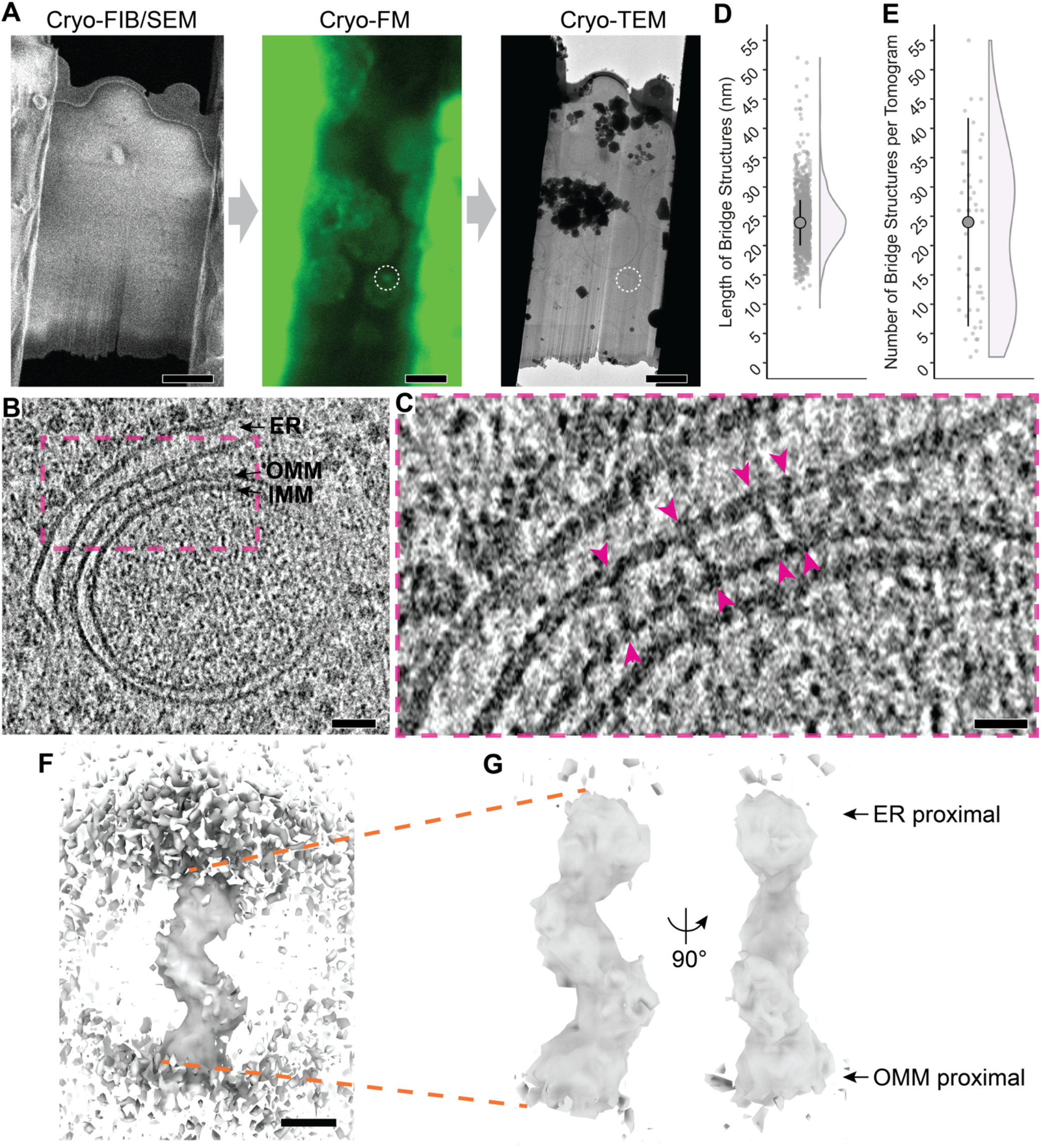
ERMES-mediated MCS consist of bridge structures connecting the two membranes. **A:** The cryo-CLEM workflow includes three microscope steps. Thinning of cells into lamellae by cryo-FIB milling visualized by scanning EM (SEM); cryo-FM of lamellae to localize Mdm34-mNeonGreen marked ERMES puncta (white dashed circle); and cryo-transmission EM (TEM) for acquisition of electron cryo-tomograms. **B:** Virtual slice through an electron cryo-tomogram acquired at an Mdm34-mNeonGreen punctum, showing an ER-mitochondrial MCS. The ER, OMM and IMM are indicated. **C:** Zoom into the region in dashed pink box in B. The arrowheads indicate bridge-like connections between the ER and the OMM. **D:** The length of the bridge structures in nm. Dot plot and half-violin plot. Large dot indicates mean (24.2 nm, N=1098 bridges), lines the SD. **E:** The number of bridge structures found per electron cryo-tomogram acquired at Mdm34-mNeonGreen puncta. Dot plot and half-violin plot. Large dot indicates median (24, N=51 tomograms), lines MAD. **F:** 3D map of the bridge structure obtained by STA. The ER (top) and OMM (bottom) membranes are partially visible. **G:** STA map at higher contour level than in F, therefore membranes are not visible. Two views rotated by 90° along the major axis. The map represents the cytosolic portion of the bridge, with regions proximal to the ER and OMM indicated. Scale bars are 3 µm in A, 50 nm in B, 20 nm in C, and 5 nm in F.

Importantly, the cryo-CLEM approach provided us with confidence that the imaged MCS contained a significant amount of ERMES (Figure 2B). Within ER-mitochondria MCS we found numerous dense, bridge-like structures spanning the gap between the two organelles (Figures 2C). These structures were on average 24.2 nm in length (standard deviation (SD)=4.76 nm, N=1098 bridges, Figure 2D). A median of 24 bridge structures were observed per tomogram (median absolute deviation (MAD)=17.8, N=51 tomograms) (Figures 2E). The number of bridge structures is in close agreement with the number of molecules of Mmm1, Mdm12 and Mdm34 per diffraction limited puncta. These findings suggest that the bridge structures, which are located at puncta of Mdm34-mNeonGreen, represent the ERMES complex forming a continuous structure between the ER and mitochondrial membranes.

To gain structural details of the bridges and insight into the organization of ERMES, we used subtomogram averaging (STA) to determine an average 3D map from subvolumes containing the bridge structures (Supplementary Figures S3 and S4). Our structural map fills the distance between the ER and mitochondria, likely corresponding to the full cytosolic portion of the ERMES complex (Figure 2F). There are two bends along the map which produce a zig-zag arrangement of three segments similar in size and shape (Figure 2G, Movie S1). This overall structure suggests the assembly of three SMP domains arranged consecutively in a continuous string. The order of the arrangement is deducible from information available on the three SMP domain containing components: Mmm1 is anchored in the ER and interacts with Mdm12; Mdm34 interacts with Mdm12 and with the OMM protein Mdm10 (16, 20, 21). Thus, in our map, the segment nearest the ER is likely to contain the Mmm1 SMP domain, the central segment Mdm12, and the segment nearest the OMM Mdm34. Such an organization fits with the equimolar ratio of components determined by FM. We propose that ERMES bridges the space between the ER and mitochondria through a stable complex in which each component contributes one SMP domain, strung in a zig-zag assemblage.

*In vitro*, the Mmm1 SMP domain homo-dimerizes like SMP domains of other lipid transfer proteins and could thus mediate dimerization of ERMES bridges through an interaction near the ER membrane (13, 16, 25). Since our STA approach averaged individual bridges, we could have missed such dimers. We therefore used tomographic coordinates of bridge-membrane anchor points proximal to the ER and the OMM, as well as the centers of the bridges, to determine whether the bridges had a specific arrangement relative to each other (Figure 3A). For each anchor point, we measured the distance to its nearest neighboring anchor point. We reasoned that if Mmm1 dimerized as in the crystal structure (16), the distance between two neighboring ER anchors would be shorter than between bridge centers or OMM anchor points. If there was another oligomerization at neighboring OMM anchors, the distance between the latter could be similar to ER anchor points, and it would be shorter than between neighboring center points (Figure 3B). Our analysis revealed that for ER anchors, bridge center points and OMM anchors, the distances between nearest neighbors were equivalent (median +/-MAD: 14.8 nm +/-8.0 nm, 14.3 nm +/-7.5 nm and 14.3 nm +/-7.9 nm, respectively; Figure 3C). We conclude that an arrangement involving dimerization of Mmm1 SMP domains as observed *in vitro* does not occur in the cell.

**Figure 3.**
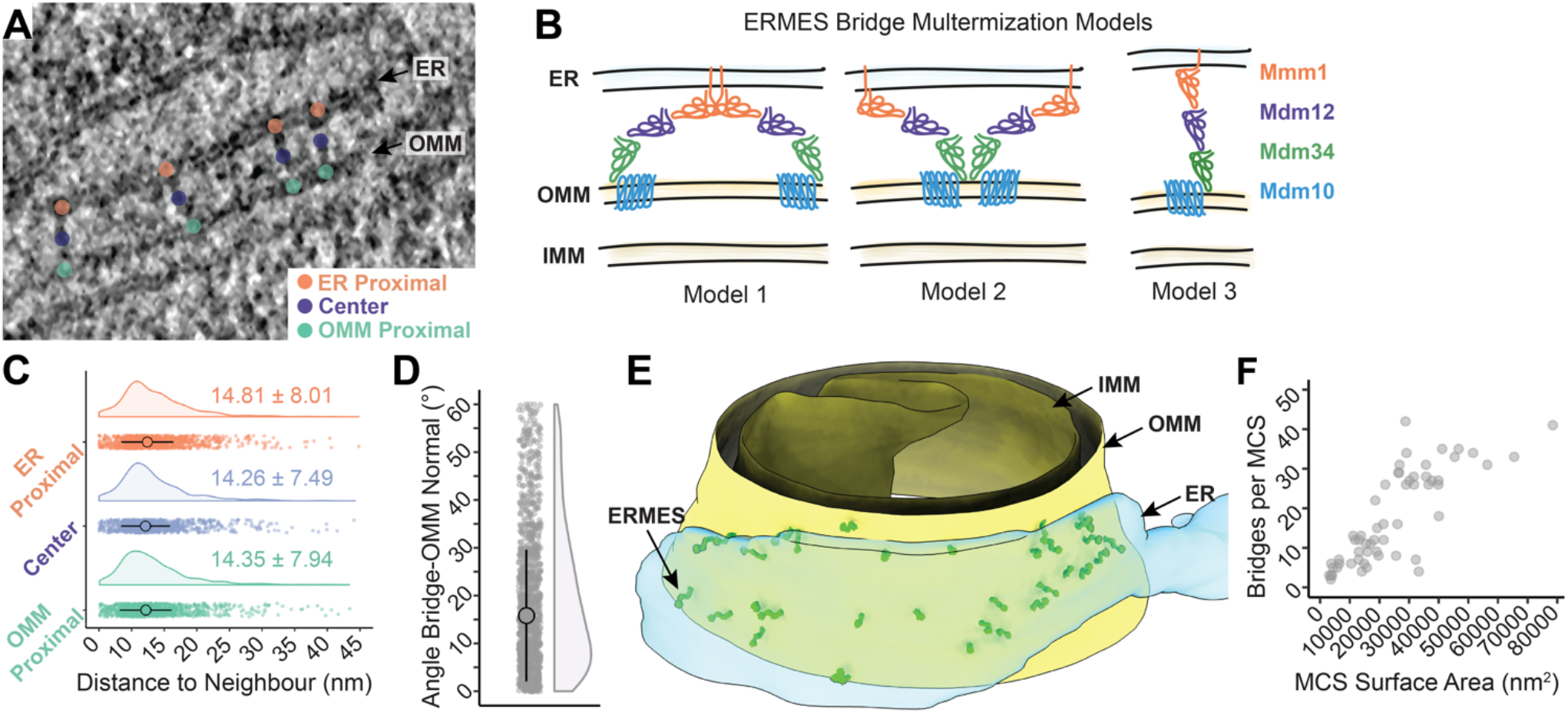
Supramolecular organization of ERMES within MCS. **A:** Coordinates of ER membrane anchor points (orange), center points (purple) and OMM anchor points (green) in electron cryo-tomograms. **B:** Three possible models how ERMES bridges could be arranged relative to each other, consistent with the STA map. Model 1: ERMES dimerizes via Mmm1 as observed *in vitro* (16). ER anchor points of neighboring bridges would be close to each other. Model 2: ERMES dimerizes via Mdm34 or Mdm10. OMM anchor points of neighboring bridges would be close to each other. Model 3: Neither model 1 nor 2 applies if neighboring bridges are similarly close at the ER, the bridge center and the OMM. **C:** Dot plot and half-violin plot of the distances between nearest neighboring bridges, measured between the ER anchor points (orange), the bridge centers (purple), and the OMM anchor points (green), respectively. Large dots indicate medians, lines MAD, both also given as numerical values. **D:** The angle by which each bridge is tilted relative to the OMM normal. Dot plot and half-violin plot. Large dot indicates median, lines MAD (N=1098 bridges). **E:** Segmentation model of an electron cryo-tomogram, showing the distribution of ERMES bridge structures within MCS. The STA map was placed at the positions of individual bridge structures, indicated by ‘ERMES’. OMM, IMM and ER are also indicated. **F:** The number of bridges per MCS, plotted as a function of the surface area of the ER membrane in contact with the OMM.

We did not include the membranes in our STA procedure (Supplementary Figure S3) to avoid these large structures dominating the alignment of subvolumes containing the bridges. We noticed, however, that many bridges were not perpendicular to the membranes, but displayed an angled orientation. To assess this observation quantitatively, we performed another alignment of the previously aligned bridge subvolumes, this time using the OMM as our target feature (Supplementary Figure S5). The difference in the angle relative to the membranes between the initial bridge alignment and the OMM alignment position was 15.8° +/-13.8° (median +/-MAD, Figure 3D). These results indicate that there is flexibility in the positioning of ERMES bridges relative to the membranes; possibly provided by hinge-like regions at the interfaces between the cytosolic and membrane parts of ERMES. This flexibility in orientation could help ERMES bridges to accommodate varying membrane curvatures of the ER and the OMM as well as potential pushing and pulling forces acting on the organelles.

When we placed the STA map back into tomograms at positions of the individual subvolumes, we observed that the bridges were distributed within the MCS in clusters of varying density (Figure 3E, Movies S2-S7). To estimate the area of membrane serviced by an individual ERMES bridge we measured the surface area of the ER membrane interfacing with the OMM. We found that one ERMES bridge occupied 1359 nm^2^ +/-483 nm^2^ (median +/-MAD, N=60 MCS) of ER membrane (Supplementary Figure S6). Thus, if equally distributed within MCS, the distance between bridges would be approximately 40 nm. However, the closest neighbor of each bridge was 15 nm away (Figure 3C), indicating that ERMES bridges are not distributed homogenously, but rather in irregular clusters. The number of bridges found per MCS correlated with the ER membrane surface area in contact with mitochondria (Figure 3F), suggesting that the number of ERMES bridges is limited to approximately 12 bridges per 10,000 nm^2^ of ER surface area. Thus, while the distribution of bridges within the MCS appears to adopt no regular pattern and the morphologies of MCS are diverse, the number of ERMES bridges per surface area (Figure 3F) and per MCS (Figure 1C) appear to be constrained, indicative of spatial regulation of ERMES organization across MCS.

We next set out to investigate the molecular architecture of the ERMES complex. Our STA map suggests a zig-zag arrangement of three SMP domains (Figure 2G). We sought to test if such an arrangement is compatible with the structural properties of the components. We predicted the structures of heterodimers (Mmm1-Mdm12, Mdm12-Mdm34 and Mdm34-Mdm10) using FoldDock (FD) (26), an AlphaFold (AF)-based tool (27), and assembled the complex based on the sequential order of the components derived from our STA map and previous findings (16, 20, 21)(Figure 4A). FD predicted a sequential tail-to-head arrangement of the SMP domains of Mmm1, Mdm12 and Mdm34, with the head-terminal loop of one subunit interacting with the crevice formed by the next subunit (Figure 4B), similar to the interfaces observed in the X-ray structure of the Mmm1-Mdm12 dimer (16).

**Figure 4.**
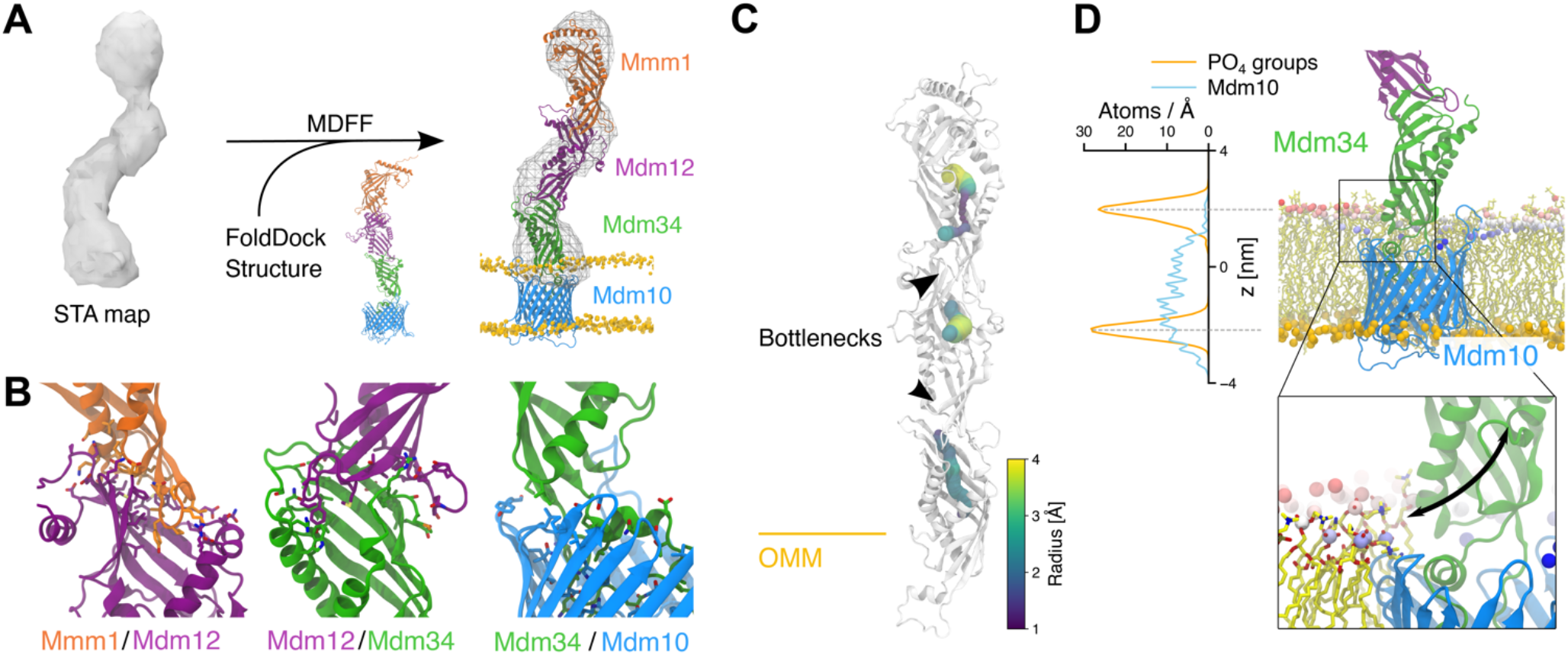
Integrative modeling of the ERMES complex. **A:** FD and MDFF approach (26, 62). The STA map was used to bias the conformation of the three SMP domains of ERMES. **B:** Predicted interfaces between Mmm1-Mdm12 (left), Mdm12-Mdm34 (center), and Mdm34-Mdm10 (right) in the model fit to the STA map. Interfacing residues (<3 Å) are shown as sticks. **C:** Radius of the cavity in the ERMES model. The three subunits show three distinct cavities, with bottlenecks at the subunit interfaces. **D:** Hydrophobic mismatch of Mdm10 (cyan) embedded in an OMM-like bilayer. Phosphate groups of the upper leaflet are colored according to their position along the z-axis (range: 10-25 Å from the bilayer center). Left: Density profiles of the Mdm10 backbone (cyan) and membrane phosphate groups (orange). Inset: Close-up representation of the Mdm34-Mdm10 interface, with the phosphate groups (orange) located near the Mdm34 cavity, indicated by black arrow.

Notably, rather than the zig-zag arrangement we observed *in situ*, the *in silico* FD prediction suggested a more ‘linear’ conformation of the complex, similar to the *in vitro* arrangement (16). To adjust the predicted complex structure to our experimental STA map, we performed atomistic Molecular Dynamics Flexible Fitting (MDFF) of the ERMES complex with an explicit lipid bilayer mimicking the OMM (Figure 4A, Supplementary Figure S7 and Movie S8). Of note, even though the STA map does not provide secondary structure information, limiting the overall accuracy of our model, fitting to the STA map only moderately modifies the inter-subunit interfaces, which remain nearly in their initial FD-predicted conformations (Figures 4A and B). Furthermore, our model does not completely occupy the STA map density around Mdm12 and Mdm34 (Figure 4A). The extra density in the map could be explained by the intrinsically disordered or dynamic protein regions that we excluded from modeling due to their poorly predicted structures, or binding of auxiliary factors such as the regulatory proteins Gem1 (21, 28), Tom7 (20) or Emr1 (29).

We next investigated whether the proposed complex is compatible with a tunnel-like lipid transport mechanism. In such a scenario, lipids should slide through the SMP domains and, importantly, move between subunits via their interfaces. In our final MDFF model, the cavities of the SMP domains do not form a continuous conduit, as they are too narrow for lipid transport at the Mdm12-Mdm34 and Mmm1-Mdm12 interfaces (Figures 4C and Supplementary Figure S8). This observation suggests that unlike other tunnel-like lipid transfer proteins such as the VPS13 family, which form large unrestrained conduits (30, 31), the lipid pathway in ERMES could be restrained by bottlenecks at the interfaces between domains. Furthermore, in our model, the cavity of Mdm34 is near the OMM surface, and the predicted interaction between Mdm34 and the OMM protein Mdm10 occurs close to residues previously found to drive Mdm34-Mdm10 complex formation (20) (Figure 4D). Notably, our simulations suggest that Mdm10 displays a hydrophobic mismatch on the cytosolic leaflet of the OMM, which therefore deforms the bilayer around Mdm10 (Figure 4D). The arrangement of the Mdm34 cavity and of the membrane around Mdm10 could facilitate the uptake and release of lipid molecules (Figure 4D, inset).

In summary, combining quantitative FM, *in situ* structural biology and integrative modeling allowed us to seamlessly build a comprehensive and highly resolved model of ER-mitochondria MCS in yeast. In this model, ERMES distributes within MCS as a discrete, two-membranes spanning complex of approximately 188 kDa in mass, with the three ERMES SMP domains arranged into a continuous structure between the two organelles. This organization could represent a controllable trajectory for lipids that differs from the shuttle or tunnel modes identified for other lipid transfer proteins (1). The exact mechanism regulating lipid passage through ERMES remains to be determined, but such a mechanism might facilitate specificity or directionality of lipid transport while being compatible with ERMES as an interorganelle tether.

## Materials and Methods

### Yeast cell culture and genetics

Liquid *Saccharomyces cerevisiae* (budding yeast) cultures were grown in synthetic complete medium without tryptophan with 2% glucose (SC-Trp) at 25°C for all experiments. Genetic tagging with either EGFP, mNeonGreen or mCherry was done according to (32), using plasmids pFA6a-EGFP-HIS3MX6, pFA6a-mCherry-KanMX and pFA6a-mNeonGreen-HIS3MX6 (32, 33).

### Yeast strains used for live cell fluorescence microscopy

Mdm34-mNeonGreen, Tom20-mCherry (WKY0401): MATa, his3Δ200, leu2-3,112, ura3-52, lys2-801, MDM34-mNEONGREEN::HIS3MX6, TOM20-mCHERRY::kanMX4

Nuf2-EGFP (WKY337): MATα, his3Δ200, leu2-3,112, ura3-52, lys2-801, NUF2-EGFP::HIS3MX6

Cse4-EGFP, Tom20-Cherry (WKY421): MATα, his3Δ200, leu2-3,112, ura3-52, lys2-801, CSE4-EGFP::HIS3MX6, TOM20-mCHERRY::kanMX4

Mdm12-EGFP (WKY0119): MATα, his3Δ200, leu2-3,112, ura3-52, lys2-801, MDM12-EGFP:: HIS3MX6

Mdm34-EGFP (WKY0432): MATα, his3Δ200, leu2-3,112, ura3-52, lys2-801, MDM34-EGFP:: HIS3MX6

Mmm1-EGFP (WKY0433): MATα, his3Δ200, leu2-3,112, ura3-52, lys2-801, MMM1-EGFP:: HIS3MX6

### Yeast strain used for cryo-CLEM /cryo-ET

Mdm34-mNeonGreen, Dnm1-mCherry (WKY0400): MATa, his3Δ200, leu2-3,112, ura3-52, lys2-801, MDM34-mNEONGREEN::HIS3MX6, DNM1-mCHERRY::kanMX4

### Quantification of the number of ERMES components per MCS

Quantification of the number of molecules of ERMES components per diffraction limited puncta was done as in (34), according to the protocol described by (19). Yeast cells expressing Cse4-EGFP Tom20-mCherry were mixed with cells expressing either Mmm1-EGFP, Mdm12-EGFP or Mdm34-EGFP, and then mounted on Concanavalin A coated 1.5 glass coverslips. The Tom20-mCherry signal was used to discriminate cells of the two strains. Stacked images (21 images over 4.2 μm z-height) were collected using the central 1024 × 1024 px of a QE80 Hamamatsu camera mounted on a Nikon Ti2 microscope equipped with Nikon Plan Apo VC 100x/1.40 NA oil objective. A NIJI LED light source with 470 nm and 550 nm LEDs was used for excitation with the filter sets 49002-ET-EGFP (FITC/Cy2) with ET470/40x, T495lpxr and ET525/50m for green fluorescence and 49005-ET-DSRed (TRITX/Cy3) with ET545/30x, T570lp and ET620/60m for red fluorescence. Stacked images were collected of green fluorescence before red fluorescence imaging. Z-stacked images of individual cells (either expressing one of the ERMES components or Cse4-EGFP Tom20-mCherry cells in anaphase-telophase) were cropped manually using Fiji (35) and then subjected to the SpotQuant analysis pipeline, which includes background subtraction, identification and masking of fluorescent spots prior to measurement of fluorescence intensity (19). The Python3 package SpotQuant was accessed at https://github.com/apicco/spotquant. To validate the workflow, the ratio of Nuf2-EGFP molecules with respect to Cse4-EGFP molecules was measured and found to be 3.5:1, in close agreement with the published ratio of 3.6:1 (36, 37).

### Cryo-FIB milling of yeast cells, cryo-CLEM and cryo-ET

Log-phase cultures of yeast expressing Mdm34-mNeonGreen and Dnm1-mCherry were pelleted and resuspended in SC-Trp with 15% dextran (w/v, Sigma 40 kDa Mw) before applying to glow discharged Quantifoil R2/2 Cu 200 mesh grids, manually back blotting and plunging into liquid ethane using a manual plunger with temperature control (38). Lamellae of vitrified yeast cells were prepared as described by (17, 39) using a Scios DualBeam FIB/SEM microscope (FEI) equipped with a Quorum PP3010T cryo-FIB/SEM preparation system. To improve lamellae stability, micro-expansion joints were made (40). Cryo-FM imaging of lamellae was done on a Zeiss Axio Imager M2m microscope equipped with a Linkam CMS 196 cryo-stage and an Axiocam 503 mono CCD camera (Zeiss). Bright field images of the grid were collected using an EC Plan-Neofluar 10x/0.30 (Zeiss) objective lens. High-magnification fluorescence images of lamellae were collected with a LD EC Epiplan-Neofluar 100x/0.75 DIC M27 (Zeiss) objective lens. A Zeiss Colibri 7 LED light source with 475 nm and 555 nm LEDs was used for excitation with the filter sets 38 HE eGFP (Zeiss) for green fluorescence (Mdm34-mNeonGreen) and 63 HE mRFP (Zeiss) for red fluorescence (Dnm1-mCherry). Room humidity was kept below approx. 25% during cryo-FM. Lamellae which contained Mdm34-mNeonGreen signals were used in the next steps of cryo-ET. Cryo-ET tilt series were collected on a Titan Krios microscope (Thermo Fisher) operated at 300 kV using a Quantum energy filter (slit width 20 eV) and a K3 direct electron detector (Gatan). SerialEM (41) was used for acquiring montaged maps of the lamellae and for tilt series acquisition. Low magnification maps (39.39 Å/px) were correlated with cryo-FM images by 2D rigid fitting using the Icy plug-in ecCLEM (42). As landmarks for registering cryo-FM and cryo-EM images, features of the lamella and yeast cell outlines, as well as holes and imperfections in the surrounding carbon support film which were visible in both cryo-FM and cryo-EM images were used. In this way regions containing Mdm34-mNeonGreen puncta were identified. In a second step, the resulting composite images were correlated to medium magnification anchor maps (10.73 Å/px), using features recognisable at low and intermediate magnification. The medium magnification maps were used to navigate tilt series acquisition. Tilt series images were acquired at 1° increment over a ±56° range in groups of 4 using a dose-symmetric tilt scheme (43, 44) at a pixel size of 2.684 Å and a defocus range from -3.5 to -6.0 μm. Total exposure time was adjusted to maintain a total dose per image of 1.3 e-/Å using a dose rate of approximately 25 e-/px/s. Exposures were fractionated as four frames acquired as LZW-compressed TIFF images without normalisation and without binning.

### Tomogram Reconstruction and Subtomogram Averaging

Frames were gain corrected, aligned and dose-weighted with the preprocessing script from the subTOM package (45) which executes IMOD’s alignframes and ctfplotter functions (46). Tilt series were aligned using IMOD’s etomo with patch tracking and 2D CTF correction applied by phase-flipping in IMOD. Tomograms were reconstructed at bin 2 using etomo by either weighted back-projection for subtomogram averaging or simultaneous iterative reconstruction technique (SIRT) for tomograms used for manual coordinate picking of the bridges’ ER and OMM anchor points, segmentations and for figures. Bridge structures were manually picked using Dynamo (47) as dipole models of ER and OMM anchor points. The central point and axis between ER and OMM anchor points were used to define the coordinates for subtomogram extraction using the subTOM package which utilizes MATLAB (MathWorks) functions adapted from TOM (48), AV3 (49, 50), and Dynamo (47). The presented STA consists of 1098 subtomograms, extracted from 51 tomograms acquired at regions of lamella corresponding to ERMES puncta imaged by cryo-FM. See Supplementary Figure S3 for a schematic of the STA procedure. For initial template generation, a random spin rotation was applied to each subtomogram to minimise the effect of the preferential missing wedge upon subtomogram alignment. Subtomogram alignment was inspected between alignment iterations using the UCSF Chimera (51) plug-in Place Object (52). Subtomograms which aligned poorly and instead ‘migrated’ to align to the ER or OMM membranes, were found by visually inspecting the position of the average STA map placed at the positions of the subvolumes within each tomogram. These misaligned subtomograms were removed from the list of subtomogram positions used for STA before the subTOM analysis was re-run. The resolution of the resulting STA map was estimated to be 29 Å from the Fourier Shell Correlation curve at 0.143, generated according to (53), implemented through subTOM (45)(Supplementary Figure S4A). For this analysis, the STA map was masked (Supplementary Figure S4B).

### Analysis of the angle between bridge and OMM

STA was used to align the subtomograms composing the bridge STA map to the OMM. The zenith rotational angle was compared between these two alignments to obtain the angle between the z-axis of the bridge STA map relative to the normal of the OMM. For this alignment, the subtomograms aligned before (Supplementary Figure S4) were repositioned so that the OMM-proximal end of the bridge was centred at the pivot point of the STA alignment search. The subtomograms were masked to include the OMM and a part of the bridge (Supplementary Figure S5C). The alignment search consisted of 5° zenith tilt-steps, from 0-60°, without in-plane rotation. The differences in zenith rotation of each subtomogram before and after this alignment were calculated in MATLAB (Mathworks) analyzed in R.

### Nearest neighbor analysis

For the nearest neighbour analysis, the Euclidean distances between all the centers of subtomograms were calculated with MATLAB (Mathworks), and for each subtomogram the minimal distance was identified as the one to the nearest neighbor. For determining the distance to the nearest neighbor of the ER and OMM anchor points, subtomograms were first re-centered on the ER or OMM anchor points, respectively, and the Euclidean distances to the nearest neighbors were determined as for the centers.

### Numerical data analysis

Data was analyzed using MATLAB (Mathworks) and R (R Core Team, 2021). For figure panels, the data was plotted using ggplot2 (54) and raincloud plots (55).

### Segmentation and ultrastructure analysis

Segmentation models of mitochondrial membranes, ER and peroxisomes were drawn manually in IMOD (46) and rendered for figure preparation in UCSF ChimeraX (56). To determine the surface area of the contact site, the ER membrane in proximity to the OMM was segmented in IMOD and its surface area was measured using imodinfo (46). The UCSF Chimera (51) plug-in Place Object (52) was used to place back the average map into tomographic coordinates and determine the number of ERMES bridges per surface area. For display of the STA average, the map was rendered in UCSF ChimeraX (56).

### Structural model and fitting

For structure prediction, the sequences were modified as follows. *Mmm1*: Truncated the ER-interacting N-terminal side after the helical segment (residues: 1-161), as in the crystal structure (16)(PDB ID:5YK6). *Mdm12*: based on the structure from (25), the used sequence comprises only part of the loop between residues 73 and 115, adding 5 residues to each side of the broken loop (73-78) and (110-115). The loop has been shown to not be necessary for function (25). *Mdm34*: Only residues 1-200 were used, corresponding to the SMP domain. The C-terminal part is expected to be partially disordered based on secondary structure prediction and its interaction with the rest of the complex is currently unknown. *Mdm10*: Three regions facing the intermembrane space and one facing the cytoplasmic side of the membrane are likely disordered or partially disordered and were not included in the used sequence (residues 95-127, 219-224, 323-406, 452-473). None of the removed loop regions is expected to strongly affect the overall architecture of the model or is interacting with other modeled subunits (25, 57).

The ERMES complex structure was predicted by combining three dimer predictions (Mmm1/Mdm12, Mdm12/Mdm34, Mdm34/Mdm10) using the FoldDock (FD) protocol (26). We combined the predictions by aligning the common subunits of the predictions. To account for the different conformations adopted by Mdm34 in the Mdm12/Mdm34 and Mdm34/Mdm10 predictions, we performed a Targeted Molecular Dynamics simulation and changed the conformation of part of the subunit to match both interfaces. The simulation was performed using Mdm34 from the Mdm10/Mdm34 predicted structure, using implicit Generalized Born solvent (e=80), added secondary structure restraints, and maintaining the regions interfacing with Mdm10 restrained to their initial positions with a harmonic force constant of 2 kcal mol^-1^ Å^-2^ on the residues 1-31, 75-130, 147-152. The simulation was performed using Charmm36m force-field parameters (58) Langevin dynamics at *T* = 310 K and a time step of 1 fs. The combination of the predictions from FD by aligning the central subunits (e.g. Mmm1/Mdm12-Mdm12/Mdm34) results in a roughly linear arrangement of subunits. In addition to FD, we performed structural prediction using Alphafold-Multimer (27, 59). The dimeric structures are predicted in the same head-to-tail conformation, yielding an overall similar architecture. Interestingly, the FD structure shows a larger aperture at the Mdm12/Mdm34 interface (Supplementary Figure S8), possibly highlighting conformational changes that could promote lipid transport.

A total of 4 POPE lipids were included in the Mmm1/Mdm12/Mdm34 complex (2:1:1) using lipid structures from the resolved crystal structures (where available) or added by similarity with other subunits. The complex (through Mdm10) was embedded in a POPC:POPE:POPI (50:35:15) membrane resembling an Outer Mitochondrial Membrane (OMM)-like composition (60), and solvated with TIP3P water. Sodium and chloride ions were added to give an ionic strength of approximately 120 mM. The total system size comprised approximately 330,000 atoms. The membrane system was prepared using Charmm-GUI (61).

All simulations were performed using Charmm36m force-field parameters and an *NPT* ensemble with *P* = 1 atm, and Langevin dynamics at *T* = 310 K. The time step was set to 1 fs, and long-range electrostatics treated by the Particle Mesh Ewald (PME) method.

After initial equilibration of the membrane (30 ns) while maintaining the protein restrained to its initial position, we biased the structure obtained with FD using a Molecular Dynamics Flexible Fitting approach (MDFF) (62) and extended the simulation to 120 ns. The densities obtained from the experiments were used to generate a grid potential with a voxel size of 5.368 Å derived from the STA map. The density was trimmed and centered to contain only the region of interest. Additionally, we partially reduced the noise of the map by applying a gaussian filter. The grid-based potential was applied in the simulation to Mmm1, Mdm12, and the solvent-exposed region of Mdm34. We performed multiple tests to choose a suitable biasing potential (gscale 0.1/0.05/0.02 kcal mol^-1^, Supplemental Figure S7). The scaling factor (g_scale) determines the weight of the experimental STA map on the total molecular potential. We find that a value of 0.02 kcal mol^-1^ consolidates the accuracy of the density and the conformational changes induced by the application of the external potential.

All the simulations were performed using NAMD Git2021-11-23 with CUDA acceleration (63). Analyses and system preparation were performed using VMD (64) and ChimeraX (58). Cavity detection was performed using CAVER 3.0 (65) with standard parameters set.

## Acknowledgements

We thank the MRC LMB facilities for light microscopy, electron microscopy and scientific computing for support with data collection and processing. We thank Marko Kaksonen for encouragement and helpful discussions. M.R.W. was supported by the Natural Sciences and Engineering Research Council (NSERC) of Canada (PGSD). This work was supported by the Medical Research Council, as part of United Kingdom Research and Innovation (also known as UK Research and Innovation) under awards MC_UP_1201/08 to W.K. and MC_UP_1201/10 to E.A.M. For the purpose of open access, the authors have applied a CC BY public copyright licence to any Author Accepted Manuscript version arising. Work in the group of W.K. was also supported by the NCCR TransCure, a National Center of Competence in Research of the Swiss National Science Foundation (SNSF) (185544). Work in the group of S.V. was supported by the SNSF (PP00P3_194807) and by the European Research Council under the European Union’s Horizon 2020 research and innovation program (grant agreement no. 803952). This work was supported by grants from the Swiss National Supercomputing Centre under project ID s1132.

## Author contributions

M.R.W. collected all experimental data. A.D.L. performed the computational integrative modeling, supervised by S.V. A.D.L. and S.V. analyzed the integrative modeling data. D.R.M. and M.R.W. developed the STA alignment procedure. M.R.W. performed STA with help by D.R.M. A.P. and M.R.W. implemented the quantitative live-cell imaging pipeline and analyzed the resulting data. M.R.W., W.K. and L.E.M. analyzed cryo-ET data and conceived the experimental strategy of the project. P.C.H. participated in yeast strain generation, cryo-FIB milling and cryo-ET data acquisition. W.K. conceived and supervised the project. M.R.W. and W.K. wrote the paper, with direct help by E.A.M., A.D.L. and S.V., as well as input from all authors.

## Notes

### Competing Interest Statement

The authors have declared no competing interest.

