## Supplementary Figures for "Supramolecular architecture of the ER-mitochondria encounter structure in its native environment"

__Supplementary Figures and Legends__

**Supplementary Figure S1**

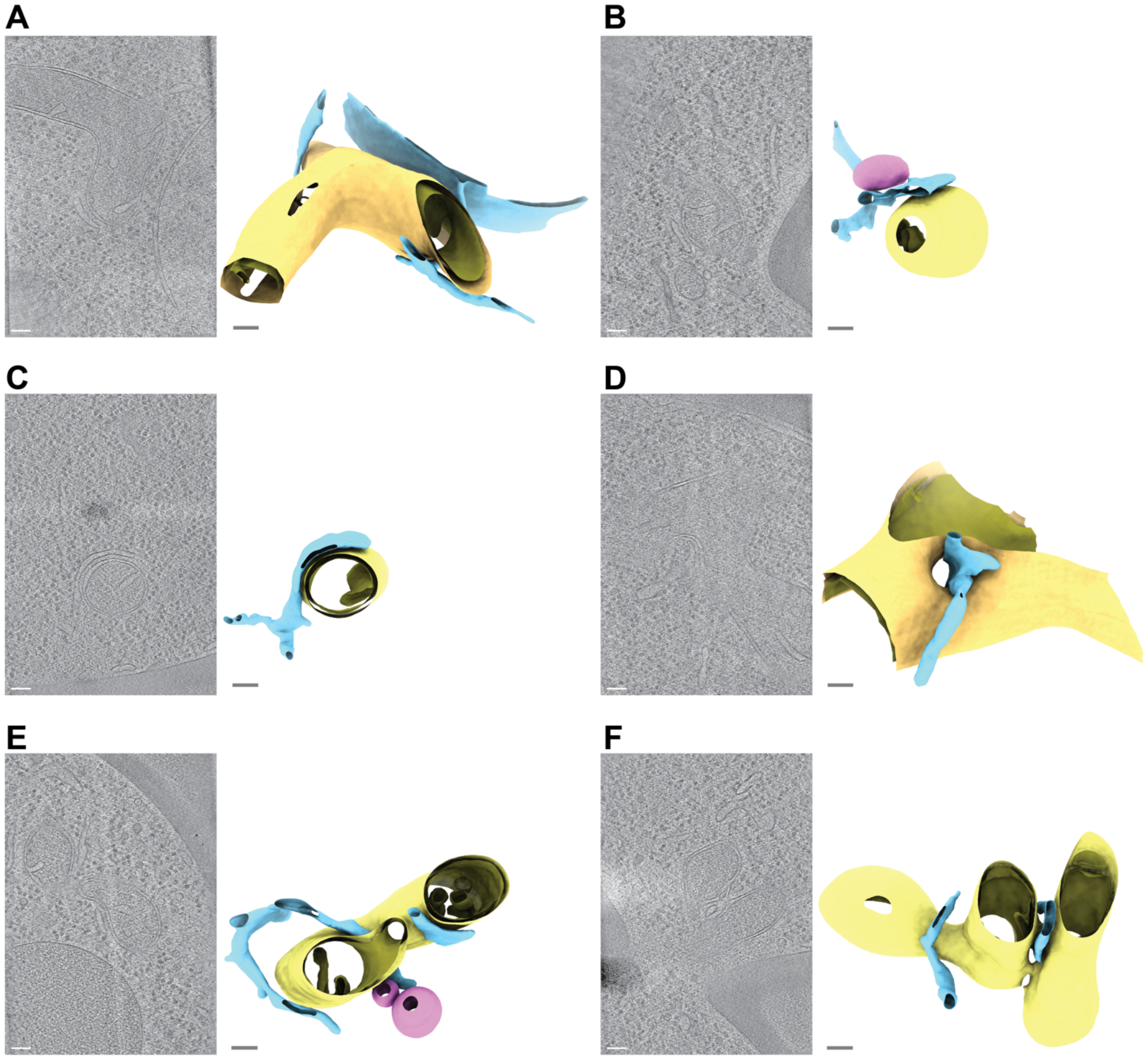


**Supplementary Figure S1. A-F:** Diversity of membrane morphologies of ER-mitochondria MCS imaged by cryo-ET at locations of Mdm34-mNeonGreen. Six representative examples from a data set of 51 tomograms are shown. Left panels are virtual slices through electron cryo-tomograms. Right panels are segmentation models of the ER (blue) the OMM (yellow), the IMM (dark yellow) and peroxisomes (pink). The segmentation models are rotated relative to the virtual slices to better visualize the MCS. The example in C is the same as shown in Figure 3E. Scale bars are 100 nm.

**Supplementary Figure S2**

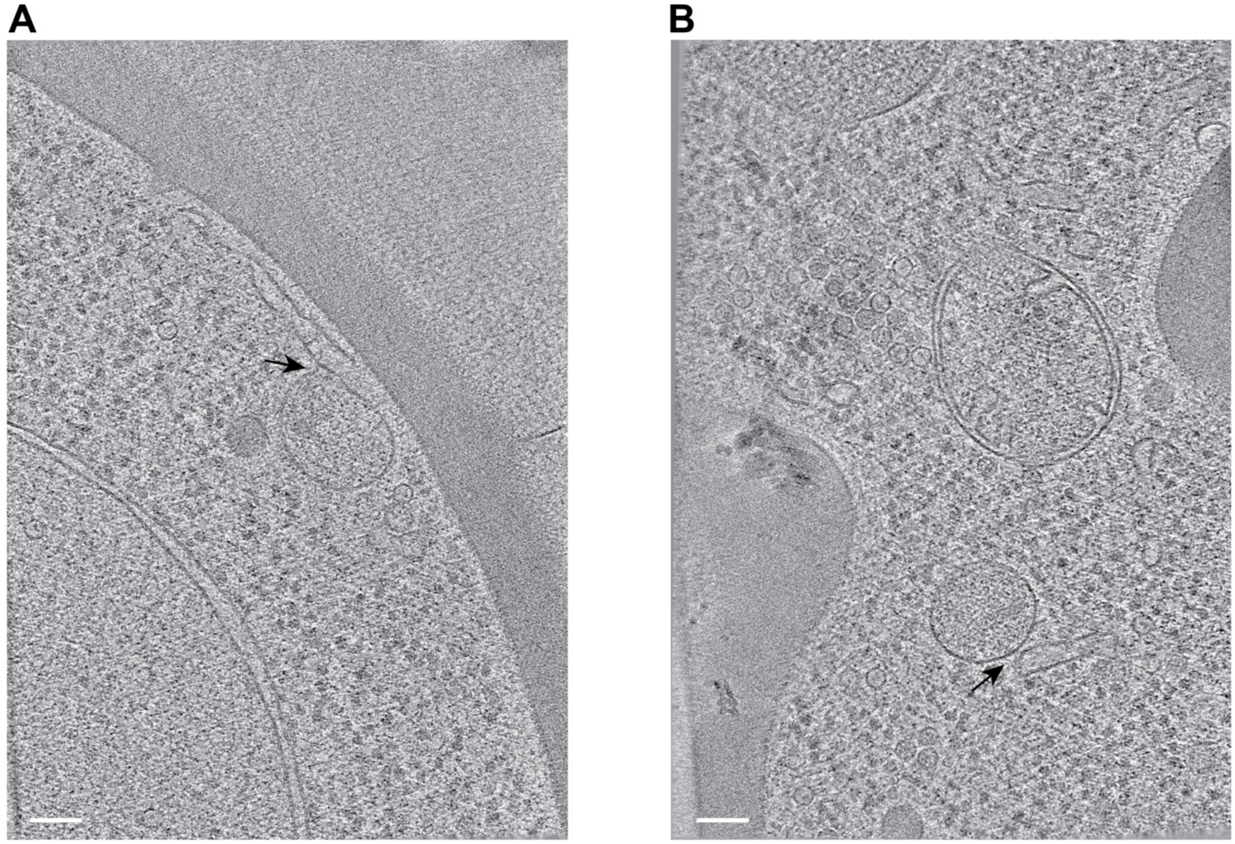


**Supplementary Figure S2. A and B:** ER-peroxisome contacts imaged by cryo-ET at locations of Mdm34-mNeonGreen. Approximately 15% of Mdm34-mNeonGreen puncta contained ER-peroxisome MCS (indicated by black arrows) rather than ER-mitochondria MCS. Peroxisomes are identified as spheroid single-membrane vesicles more than 100 nm in diameter with dense interior (1, 2). Scale bars are 100 nm.

### Supplementary Figure S3

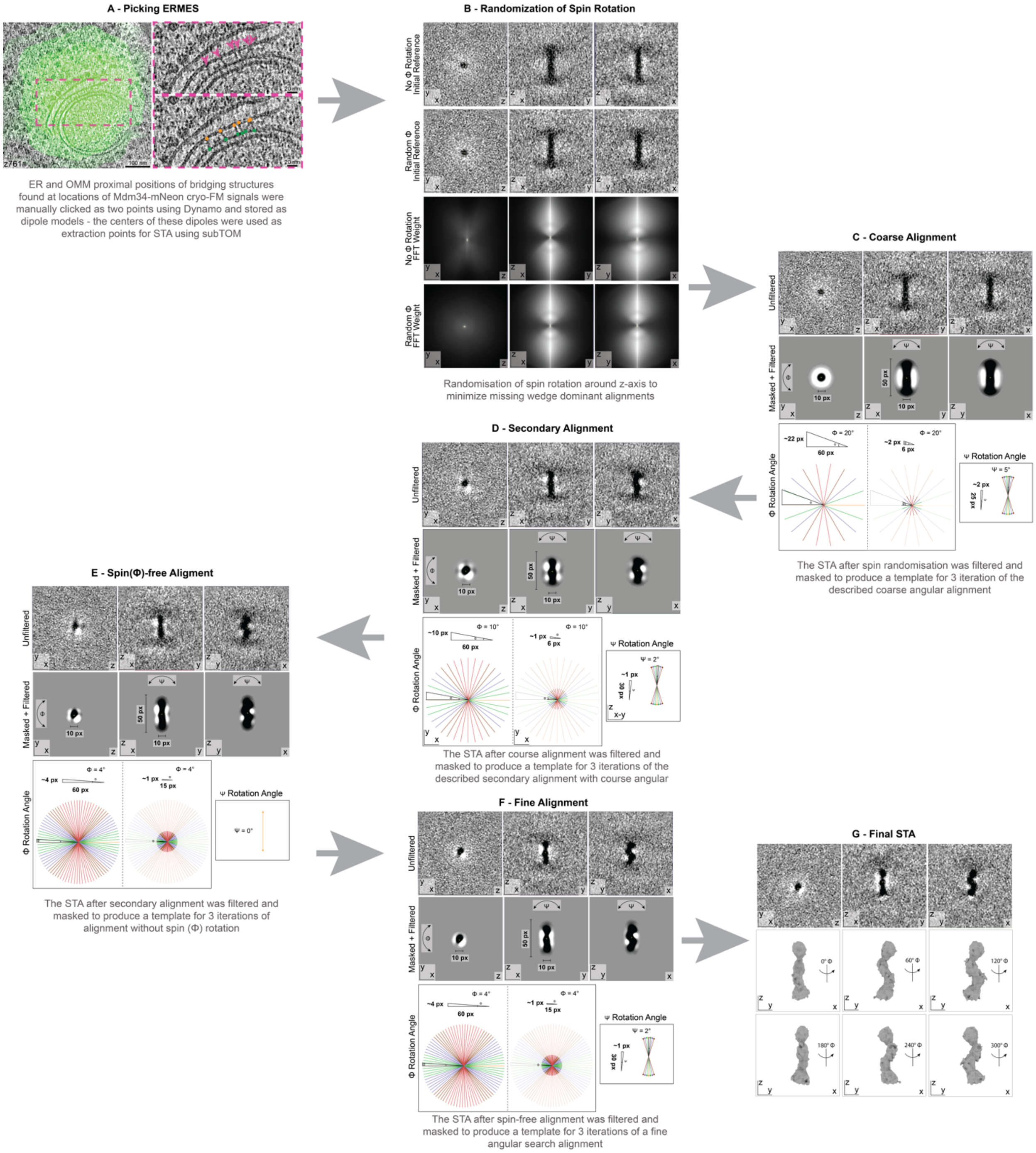


**Supplementary Figure S3.** Subtomogram averaging pipeline. The strategy using sequential alignment steps is outlined. The data set consisted of manually picked coordinates of the ER and OMM anchor points of 1098 bridge structures from 51 electron cryo-tomograms at positions of Mdm34-mNeonGreen signals. Images in A are the same as shown in Figure 2B and C, with modifications to the overlay.

**Supplementary Figure S4**

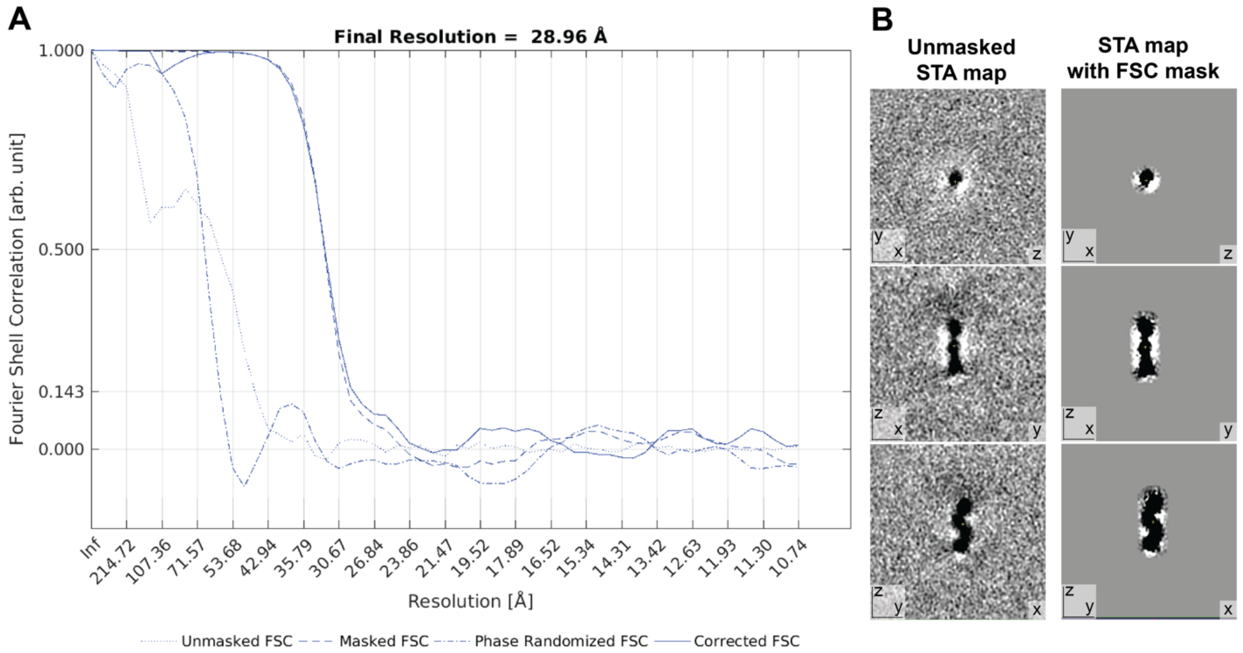


**Supplementary Figure S4. A:** Fourier shell correlation and resolution estimate, calculated according to (3). The final resolution corresponds to FSC = 0.143. **B:** Left panel: Three virtual slices through the unmasked STA map. Right panel: Three virtual slices through the STA map masked for FCS calculation.

**Supplementary Figure S5**

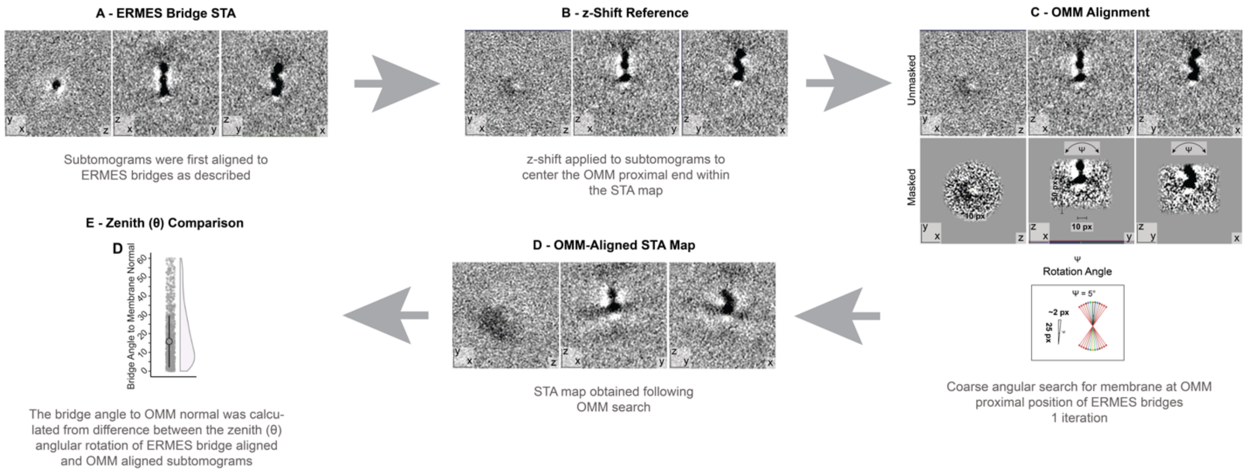


**Supplementary Figure S5.** STA alignment strategy to determine the angle between the bridges and the OMM. The graph in E is the same as shown in Figure 3D.

**Supplementary Figure S6**

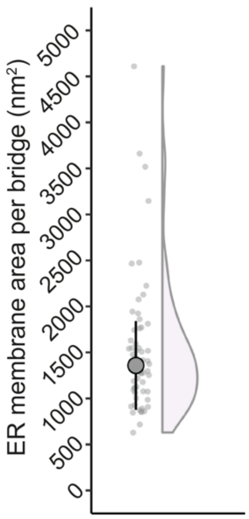


**Supplementary Figure S6.** Dot plot and half-violin plot of the surface area of the ER membrane serviced by one ERMES bridge. Large dot indicates median, lines indicate median absolute deviation.

**Supplementary Figure S7**

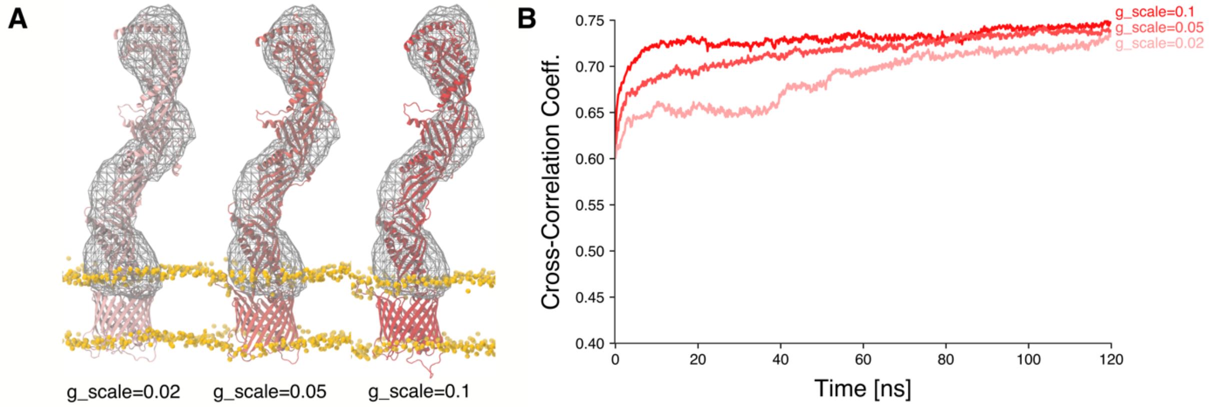


**Supplementary Figure S7.** Molecular Dynamics Flexible Fitting (MDFF) protocol. **A:** Final conformations of ERMES obtained from the MDFF simulations after fitting with different scaling factors. The scaling factor (g_scale) determines the weight of the experimental STA map on the total molecular potential. **B:** Cross-correlation coefficient calculated from the simulated (filtered to 30 Å resolution) and the experimental map.

**Supplementary Figure S8**

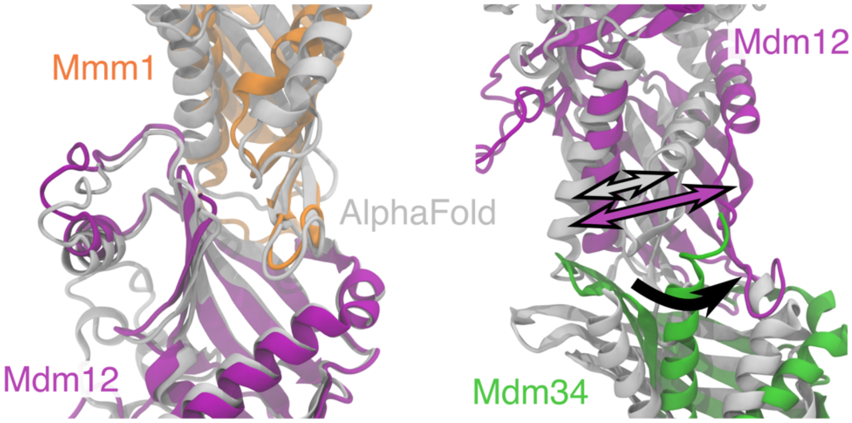


**Supplementary Figure S8.** Differences between FD (in color) and AF multimer (gray) predicted structures. While initial predictions of the complex structure using AF multimer (4, 5) and FD (6) yield a nearly identical Mmm1-Mdm12 interface, the Mdm12-Mdm34 FD-interface shows a larger aperture which tends to close upon fitting to the cryo-ET STA map. This difference between AF and FD highlights a lack of dynamic information as a current limitation of structural prediction, but could help infer possible regulatory mechanisms of ERMES function, as these regions could undergo conformational changes to promote lipid transport.

**Movie captions:**

**Movie S1:** The STA map calculated from bridge structures, in the first half of the movie at low contour level to visualize parts of the ER membrane (top) and the OMM (bottom). In the second half of the movie, the contour level is increased.

**Movie S2-S7:** Segmentation models of ER-mitochondria contact sites, imaged by cryo-ET at locations of Mdm34-mNeonGreen. The STA map was placed at the positions of individual bridge structures (shown in green). The ER is shown in cyan (semi-transparent), OMM in yellow, IMM in mustard, and peroxisomes in pink (in Movies S3 and S6 only).

**Movie S8:** Model of the ERMES complex including the OMM, obtained by integrative modeling (FD followed by MDFF into the STA map).
